# Non-linear and time-domain sleep qEEG features predict CSF protein damage markers in early Alzheimer’s disease

**DOI:** 10.1101/2025.03.28.645905

**Authors:** Anna Michela Gaeta, Lorena Gallego Viñarás, Ferran Barbé, Pablo M. Olmos, Reinald Pamplona, Farida Dakterzada, Arrate Muñoz-Barrutia, Gerard Piñol-Ripoll

**Affiliations:** Servicio de Neumología, Hospital Universitario Severo Ochoa, Leganés, Spain; Departamento de Bioingeniería, Universidad Carlos III de Madrid, Leganés, Spain; Instituto de Investigación Sanitaria Gregorio Marañón, Madrid, Spain; Group of Translational Research in Respiratory Medicine, Hospital Universitari Arnau de Vilanova and Santa Maria, IRBLleida, Lleida, Spain; Centro de Investigación Biomédica en Red de Enfermedades Respiratorias (CIBERES), Madrid, Spain; Departamento de Teoría de la Señal y las Comunicaciones, Universidad Carlos III de Madrid, Leganés, Spain; Department of Experimental Medicine, University of Lleida-Biomedical Research Institute of Lleida (UdL-IRBLleida), Lleida, Spain; Unitat Trastorns Cognitius, Clinical Neuroscience Research, IRBLleida-Santa Maria Lleida University Hospital, Lleida, Spain; Institut de Recerca Biomèdica de Lleida (IRBLleida), Lleida, Spain; Alzheimer’s Disease and Other Cognitive Disorders Unit, Neurology Service, Hospital Clínic de Barcelona, Fundació de Recerca Clínic - Institut d’Investigacions Biomèdiques August Pi i Sunyer (IDIBAPS), Universitat de Barcelona, Barcelona, Spain

**Author notes:** Corresponding author Address: Escuela Politécnica Superior, Avda. de la Universidad, 30, 28911 Leganés (Madrid) España. Anna Michela Gaeta and Lorena Gallego Viñarás contributed equally to this work.

**Keywords:** Alzheimer’s disease, sleep qEEG, CSF protein oxidation-derived markers, oxidative stress, entropy, skewness, maximum value, variance, ensemble methods

## Abstract

**Study Objectives:** In preclinical Alzheimer’s disease (AD), oxidative stress plays a key role in the pathogenesis by promoting non-enzymatic protein modifications that lead to the accumulation of oxidized species, established biomarkers of oxidative damage detectable in cerebrospinal fluid (CSF). These molecular alterations contribute to impaired proteostasis and the dysfunction of sleep-regulating neural circuits, resulting in altered sleep electroencephalographic patterns. Due to the invasiveness of CSF sampling, quantitative electroencephalography (qEEG) is proposed as a non-invasive alternative for assessing oxidatively modified protein levels via Machine Learning (ML).

**Methods:** Forty-two mild-to-moderate AD patients underwent polysomnography (PSG). QEEG features were extracted. CSF protein oxidation markers levels —glutamic semialdehyde, aminoadipic semialdehyde, Nε-carboxyethyl-lysine, Nε-carboxymethyl-lysine, and Nε-malondialdehyde-lysine —were assessed by gas chromatography/mass spectrometry, and ML models trained to predict CSF biomarker levels. Model generalizability was validated using EEG data from healthy controls.

**Results:** qEEG features from slow-wave sleep (SWS) and rapid eye movement (REM) sleep, particularly over frontal and central regions, yielded R^2^ > 0.625 for patients biomarker prediction.

**Conclusion:** qEEG is a non-invasive, scalable tool for detecting AD-related oxidative processes, with potential implications for early diagnosis and risk stratification.

**Highlights:**

- Sleep EEG features predict CSF oxidative biomarkers in early Alzheimer’s disease.
- Peak-to-peak amplitude, variance, and entropy were the top predictive features.
- NREM sleep, especially SWS, and REM sleep provided the most informative EEG signals.
- EEG-based models achieved accurate, non-invasive biomarker estimation.
- Sleep qEEG may enable early detection and risk stratification in Alzheimer’s.

## 1. Introduction

Alzheimer’s disease (AD) is the primary cause of progressive neurodegeneration, accounting for the majority of dementia cases. Its projected rise due to population aging poses a critical public health challenge, highlighting the need for early diagnostic strategies and disease-modifying therapies (Masters et al., 2015; Alzheimer’s Association, 2025). AD is marked by progressive neuronal loss in cortical and hippocampal regions, associated with extracellular amyloid-beta (Aβ) deposits and intracellular Tau aggregates forming neurofibrillary tangles (Masters et al., 2015).

In the preclinical stages of AD, pathological changes progress silently before cognitive symptoms emerge, creating a neurotoxic environment that disrupts neuronal network integrity (Qiu et al., 2009; Masters et al., 2015; Ferrer, 2023; Jové et al., 2021; Dhapola et al., 2024; Tönnies and Trushina, 2017; Wang and Michaelis, 2010). Identifying biomarkers that capture early disease alterations is crucial as timely intervention may delay or prevent progression (Tan et al., 2014; Jack Jr et al., 2024).

It is known that oxidative stress—stemming from an imbalance between pro- and antioxidants—is a major contributor to early mechanisms preceding overt AD pathology (Ferrer, 2023; Jové et al., 2021; Dhapola et al., 2024; Tönnies and Trushina, 2017; Wang and Michaelis, 2010). Oxidative stress triggers excess reactive oxygen species (ROS) production that damages key macromolecules, disrupting cellular homeostasis and fostering amyloid-β (Aβ) production and Tau hyperphosphorylation (Tönnies and Trushina, 2017)(Tönnies and Trushina, 2017).

Brain proteins are particularly vulnerable to oxidative damage, which results in irreversible structural and functional alterations (Sultana and Butterfield, 2024; Tramutola et al., 2017). This process promotes the accumulation of oxidized species, such as advanced glycation end-products (AGEs), advanced lipoxidation end-products (ALEs), and protein carbonyls—recognized biomarkers of oxidative protein damage in biological samples (Tramutola et al., 2017; Thorpe and Baynes, 2003; Requena et al., 2001; Alkhalifa et al., 2025).

Quantitative analyses of specific carbonyls, AGEs and ALEs in the cortical brain tissue of AD patients revealed a notable, though variable, elevation in the levels of aminoadipic semialdehyde (AASA), glutamic semialdehyde (GSA), Nε-malondialdehyde-lysine (MDAL), Nε-carboxymethyl-lysine (CML), and Nε-carboxyethyl-lysine (CEL) when compared to age-matched controls (Pamplona et al., 2005; Quevenco et al., 2017). Besides, CML levels in the cerebrospinal fluid (CSF) of AD patients are significantly higher compared to those of cognitively healthy individuals (Bär et al., 2003; Ahmed et al., 2005).

These findings reflect elevated oxidative damage and impaired proteostasis in AD (Pamplona et al., 2005; Quevenco et al., 2017; Sultana et al., 2009), mechanistic pathways underlying synaptic failure and neurodegeneration, which ultimately compromise the integrity of brain regions and neural networks involved in memory and sleep–wake regulation (Morrone et al., 2023; Matsumoto and Tsunematsu, 2021; Minakawa et al., 2019; Czekus et al., 2022; Tönnies and Trushina, 2017). Such pathological changes manifest as altered oscillatory patterns in sleep electroencephalogram (EEG), particularly during slow wave sleep (SWS) and rapid eye movement (REM) sleep, which are crucial for synaptic homeostasis and cognition (Czekus et al., 2022; Minakawa et al., 2019; Romanella et al., 2021; Matsumoto and Tsunematsu, 2021).

Although CSF oxidative stress biomarkers are not yet validated for clinical use, they provide valuable insight into early molecular damage and may serve as complementary indicators of biological vulnerability in individuals at risk for AD (Di Domenico et al., 2016). However, their widespread use is hindered by the invasive nature of sampling and the limited reliability of oxidized protein markers in peripheral biofluids (Haddad et al., 2019; Klimiuk et al., 2019; Kim, 2022).

In this context, sleep EEG emerges as a promising, non-invasive, and cost-effective tool for assessing early oxidative stress-related processes. By capturing specific neurophysiological alterations associated with oxidative damage and excitotoxic cascades (Ballesteros et al., 2013; Kamat et al., 2016), quantitative EEG (qEEG) overcomes the limitations of brain tissue sampling (Pamplona et al., 2005) and the impracticality of routine CSF analysis while maintaining high specificity for early AD-related changes (Di Domenico et al., 2016; Kim, 2022; Bär et al., 2003; Ahmed et al., 2005). Integrating qEEG with computational approaches, including Machine Learning (ML), could offer a powerful strategy for indirectly inferring physiological profiles, enabling early detection, risk stratification, and monitoring of AD progression.

QEEG enables automated, objective analysis of brain connectivity and network dynamics. Traditional sleep qEEG analyses, based on spectral power and sleep stage scoring, provide a limited perspective on neurophysiological changes (Thakor and Tong, 2004). In contrast, non-linear dynamics and time-domain qEEG measures offer a more comprehensive characterization of brain network complexity and functional organization (Thakor and Tong, 2004; Jeong, 2002, 2004). These advanced metrics exhibit greater sensitivity, capturing subtle preclinical neural alterations that conventional methods often overlook (Jeong, 2002).

We hypothesise that specific non-linear and time-domain features of sleep EEG act as quantitative markers of neural activity, allowing us to approximate individual profiles of oxidative stress-related biomarkers. These features may capture network vulnerabilities even before the onset of cognitive impairment or overt neuropathology. We propose that EEG-derived metrics can uncover underlying biochemical and physiological patterns characteristic of Alzheimer’s disease-associated oxidative dysregulation.

ML techniques, known for their ability to detect complex patterns and automate decision-making, are well-suited for EEG analysis. By handling high-dimensional data and capturing subtle neural signatures, ML holds significant potential for improving the diagnostic accuracy and efficiency of qEEG-derived biomarkers in AD research (Bzdok et al., 2018; Gemein et al., 2020). While the application of ML to qEEG has gained traction in AD studies (Al-Nuaimi et al., 2021; Poil et al., 2013; Gaeta et al., 2024; Gallego-Viñarás et al., 2024), these methods have not been widely employed to estimate CSF biomarker profiles related to non-enzymatic oxidative damage, especially during sleep. Filling this gap could yield critical insights into early oxidative stress-driven neurodegeneration.

This study aims to address the limitations of invasive and costly CSF biomarker collection by investigating the feasibility of noninvasive, EEG-based approaches for early oxidative risk stratification (Figure 1). By identifying electrophysiological correlates of oxidative stress, we seek to support the development of personalised, sleep-based prevention strategies for individuals at risk of AD.

**Figure 1:**
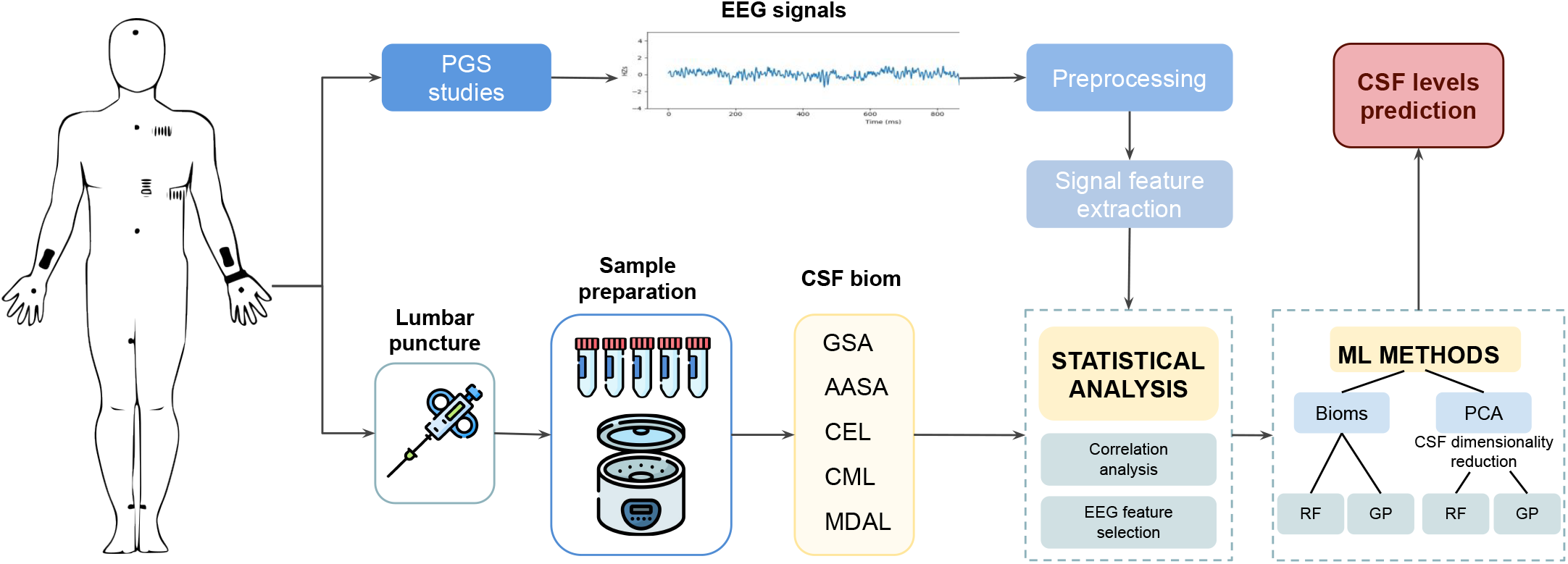
Workflow diagram illustrating the study process. EEG signals were obtained from PSG recordings and analyzed to extract temporal and nonlinear features. Simultaneously, CSF biomarker levels were measured from patient samples. Statistical analysis was performed to examine associations between EEG features and biomarker levels. Machine learning models, including Principal Component Analysis (PCA), Random Forest (RF), and Gaussian Process (GP), were then applied to predict protein levels based on EEG-derived features.

To this end, we evaluate the predictive capacity of diverse quantitative sleep EEG features in approximating individual oxidative stress biomarker profiles, including GSA, AASA, CEL, CML, and MDAL, using ML-driven modeling to uncover relevant AD pathophysiological signatures.

## 2. Materials and Methods

### 2.1. Sample and study design

This is an ancillary study from NCT02814045, a prospective trial designed to evaluate the evolution of AD patients with and without Obstructive Sleep Apnea (OSA) after one year of follow-up. The study was conducted in line with the Declaration of Helsinki and was approved by the human ethics care ethics committee of the Hospital Arnau de Villanova (CE-1218). General recruitment and screening procedures are previously described (Gaeta et al., 2020). In short, eligible patients were recruited from the Cognitive Disorders Unit at the Hospital Universitari Santa Maria (Lleida, Spain), between 2014 and 2018.

The subject involved in the analysis met the following criteria: (1) drug naïve; (2) aged over 60 years; (3) diagnosed with AD according to the NIA-AA criteria (McKhann et al., 2011); (4) diagnosed with mild-moderate cognitive impairment (an early stage of memory or other cognitive ability loss) using the Mini-Mental State Examination (MMSE ≥ 20) (Lobo et al., 1999); and 5) having signed informed consent along with the responsible caregiver (and/or legal representative if different from the responsible caregiver). Exclusion criteria for this study were: (1) history of sleep disorders; (2) comorbidities as cancer, severe depression, severe renal or hepatic insufficiency, severe cardiac or respiratory failure; (3) history of mental disorders according to DSM-V-TR™ criteria; (4) history of untreated (or treated by less than 3 months before the screening visit) vitamin B12 or folate deficiency; (5) history of untreated thyroid disease; (6) current use of medication under investigation or beta-blockers, antidepressants, neuroleptics, hypnotics use for less than 15 days before the conduction of polysomnography (PSG); (7) Magnetic Resonance Image (MRI) evidence of hydrocephalus, stroke, space-occupying lesion, or any clinical relevant central nervous system disease besides AD; (8) excessive alcohol intake (>280 gr/week); and (9) visual and/or communication disturbances affecting the compliance with the study procedures.

Eligible patients undergoing overnight PSG, blood, and CSF samples were collected to determine the Apolipoprotein E genotypes (APOE) and the levels of the biomarkers, respectively. Sociodemographic and clinical variables were collected, including age, sex, ethnicity, anthropometric data, clinical history, and sleep-related symptoms.

### 2.2. Sample collection

Fasting blood and CSF samples were obtained between 8:00 and 10:00 a.m. to minimize circadian-related fluctuations. CSF was collected in polypropylene tubes and centrifuged at 2000×g for 10 minutes at 4°C to remove cells and other insoluble components. Blood samples were drawn into EDTA tubes, and plasma was separated by centrifuging the blood at 1500 rpm for 20 minutes. The buffy coat was isolated for DNA extraction and subsequent APOE genotyping. All samples were stored at −80°C until further analysis. The samples were provided with the assistance of the IRBLleida Biobank (B.0000682) and PLATAFORMA BIOBANCOS PT17/0015/0027 and were subsequently processed for biomarker analysis.

### 2.3. APOE genotyping

APOE genotype was performed with a Maxwell^®^ RCS blood DNA kit (Promega, USA) from 2 mL of a peripheral blood sample, using 20 microliters of DNA. Participants were dichotomized according to APOE genotype as a homozygous or heterozygous carrier (APOEε4+) or non-carrier (APOEε4-) (Zhong et al., 2016).

### 2.4. AD core biomarker measurement

Commercial kits (Innotest^®^ β-Amyloid1-42; Innotest^®^ hTAU Ag; and Innotest^®^ Phospho-TAU181P, Fujirebio-Europe, Gent, Belgium) were used to evaluate the main AD biomarkers (Aβ42, t-tau, and p-tau). Analyses were performed by board-certified laboratory technicians masked to clinical data in one round of experiments using one batch of reagents. Intra-assay coefficients of variation were lower than 10%–13% for internal quality control samples (two per plate). The amounts of product were expressed as pg/ml. Concentrations of Aβ42 below 600 pg/ml, t-tau exceeding 425 pg/ml, and p-tau above 65 pg/ml were classified as positive or indicative of abnormality (Dakterzada et al., 2023).

### 2.5. Oxidative damage markers measurement

A total of five biomarkers of non-enzymatic protein modifications indicative of oxidative stress were detected and quantified, as previously described (Pamplona et al., 2005; Dakterzada et al., 2023). These included markers of protein oxidation—GSA and AASA; CEL and CML; and MDAL. All were identified as trifluoroacetic acid methyl ester (TFAME) derivatives in acid-hydrolyzed, delipidated, and reduced CSF protein samples using gas chromatography-mass spectrometry (GC/MS).

Briefly, protein samples (500 µg) were delipidated twice by adding methanol:chloroform (2:1, v/v). Following centrifugation, the methanol phase containing proteins was precipitated with cold trichloroacetic acid (TCA) and reduced with 500 mM sodium borohydride in 0.2 M borate buffer (pH 9.2) after overnight incubation at room temperature. The samples were then reprecipitated with TCA, and a mixture of deuterated internal standards (IS) for each marker was added, including [2H8]Lys (d8-Lys), [2H5]5-hydroxy-2-aminovaleric acid (d5-HAVA) as a stable derivative for GSA, [2H4]6-hydroxy-2-aminocaproic acid (d4-HACA) as a stable derivative for AASA, [2H4]CEL (d4-CEL), [2H4]CML (d4-CML), and [2H8]MDAL (d8-MDAL). Proteins were then hydrolyzed with hydrochloric acid (HCl) at 155°C and dried under vacuum.

TFAME derivatives were prepared by adding methanolic-HCl followed by trifluoroacetic anhydride incubation. The derivatized protein hydrolysates were resuspended in dichloromethane and transferred to autosampler vials for SIM-GC/MS analysis. Reference materials from NIST and quality control samples were included every five samples. Four microliters were injected in splitless mode into an Intuvo 9000 GC system equipped with an HP-5MS UI capillary column and coupled to a 5977 MSD (Agilent Technologies, Barcelona, Spain). The injector temperature was maintained at 250ºC.

The oven temperature program was as follows: 110°C for 5 min, ramped to 150°C and held for 2 min, then to 240°C for 5 min, followed by 300°C for 15 min, and held at 300°C for an additional 5 min. A 3-minute post-run at 110°C completed the cycle, yielding a total runtime of 52 min. Data acquisition was performed in selected ion monitoring (SIM) mode across an m/z range of 30–650. The MSD source temperature was set at 250°C, and the transfer line temperature at 310°C. Helium served as the carrier gas. Detected ion pairs were: Lys and d8-Lys, m/z 180 and 187, respectively; HAVA (GSA) and d5–HAVA (d5-GSA), m/z 280 and 285, respectively; HACA (AASA) and d4–HACA (d4-AASA), m/z 294 and 298, respectively; CEL and d4-CEL, m/z 379 and 383, respectively; CML and d4-CML, m/z 392 and 396, respectively; and MDAL and d8-MDAL, m/z 474 and 482, respectively. The concentration of each marker was expressed as micromoles of GSA, AASA, CEL, CML, or MDAL per mole of lysine.

### 2.6. Polysomnography

Polysomnography (PSG) was performed overnight using the Philips Respironics Alice 6 LDx system. Each recording included two bilateral EOG signals, six EEG channels from the central (C3-A2, C4-A1), frontal (F3-A2, F4-A1), and occipital (O1-A2, O2-A1) regions, as well as chin EMG, respiratory effort from chest and abdomen, airflow, pulse oximetry, ECG, body position, snoring, and leg movements.

Manual scoring and diagnosis were conducted by expert physicians in accordance with the American Academy of Sleep Medicine (AASM) guidelines, which included annotating artifact segments due to movements or poor sensor contact that hindered physiological signal interpretation (Berry et al., 2015). Sleep quality measures included total sleep time (TST), sleep efficiency (SE), and arousal index (AI). TST was calculated by subtracting sleep latency and wakefulness after sleep onset from total recording time. SE was the percentage of TST relative to time in bed (TIB), defined as the period from lights-off to lights-on. AI indicated the number of arousals per hour of sleep (Berry et al., 2015).

Additional PSG measures included time spent awake (W), which is defined as the duration and percentage of TIB spent awake. Latency to stage 1 (N1) of non-rapid eye movement (NREM) sleep was measured from lights off until the first N1 episode. Latency to REM sleep was similarly defined. N1, N2, and N3 sleep durations were quantified in minutes and as percentages of TST, with N3 also recognized as SWS. REM sleep duration was similarly measured (Berry et al., 2015).

For respiratory parameters, apnea was defined as a reduction in breathing amplitude to below 10% of baseline for over 10 seconds, while hypopnea involved a decrease in airflow by at least 30% for more than 10 seconds, leading to arousal or oxygen desaturation (a decrease in oxygen saturation greater than 3%). The apnea-hypopnea index (AHI) was calculated as the number of apnea and hypopnea events per hour of sleep. Cumulative sleep time percentage with oxygen saturation (SpO2)

< 90% was defined as CT90, and average SpO2 was the mean oxygen saturation during the study. Obstructive sleep apnea (OSA) was defined as an AHI ≥ 15 events/hour (Berry et al., 2015).

### 2.7. Healthy Controls Database

To validate model predictions and compare AD patient data with healthy individuals, we compiled a database of sleep EEG recordings from three publicly available datasets. All included subjects were free from cardiac, neurological, and psychiatric disorders and had no clinical diagnosis of sleep disorders.

- **ISRUC-SLEEP Dataset (Subgroup III):** Includes recordings from 10 healthy individuals (9 males, 1 female; mean age ~ 40 years). EEG signals were recorded from four channels (C3-A2, C4-A1, F4-A1, O2-A1), filtered between 0.3–35 Hz, and sampled at 200 Hz (Khalighi et al., 2016).
- **DREAMS Open Dataset - Healthy (DOD-H):** Contains data from 25 participants with no history of recent severe illness. EEG was recorded using a Siesta 802 device with six channels sampled at 256 Hz and filtered from 0.03–35 Hz. For this study, we used four consistent channels: C3/M2, C4/M1, F4/M1, and O2/M1 (Guillot et al., 2020).
- **Sleep Disorders Research Center (SDRC) Dataset**: Comprises recordings from 11 healthy individuals from Kermanshah, Iran. EEG was recorded at 256 Hz using a SOMNOscreen™ plus PSG system. From the 14 recorded EEG channels, we selected C4A1, C3A2, F4A1, and O2A1 for consistency (Rezaei et al., 2017).

All EEG recordings used in this study were monopolar, ensuring uniform signal referencing across datasets. Each subject’s sleep stages were scored by clinical experts, and hypnograms were available for all recordings.

Given the slight age differences across datasets, particularly between healthy controls and AD patients, we selected EEG channels from multiple brain regions—including frontal, central, and occipital sites—to minimize bias arising from regional age-related EEG variability. Notably, prior studies suggest that parietal regions may be especially sensitive to age effects in sleep EEG **?**, reinforcing our electrode selection strategy.

### 2.8. Signal preprocessing

A signal preprocessing phase is needed to homogenize the overall signal information retrieved from the different EEG recordings from AD and healthy patients and mitigate noise effects during the analysis phase, as previously performed (Bandyopadhyay and Goldstein, 2023). To preprocess and analyze signal data, Python (version 3.10.12) was employed.

The initial dataset comprised overall recorded PSG signals with annotations provided by clinical experts, following AASM guidelines as described above (Berry et al., 2015). As EEG has been considered the PSG signal of interest, the EEG signals from four channels were used: both-sided central (C3-A2, C4-A1), frontal (F4-A1), and occipital (O1-A2) signals.

EEG signals were monopolar since all of them resulted from the difference between the signal recorded in a specific brain region (C-central, F-frontal, O-occipital) and the reference electrode (A-earlobe or M-mastoid). All of them were digitally acquired with an initial sampling rate (*f s*_0_) ranging from 200 to 500 Hz.

The preprocessing phase was broken down into four primary steps:

- **Resampling**: The EEG signals were resampled for consistency across channels and subjects, addressing variations in sampling rates to an output rate (*f s* _*f*_) of 500 Hz following the AASM guidelines (Berry et al., 2015).
- **Filtering**: Bandpass filtering was applied to all signals, with cutoff frequencies set according to AASM guidelines (Berry et al., 2015). This ensured consistent spectral content within 0.3 – 35 Hz while preserving essential EEG frequencies. Specifically, frequency cuts removed the DC signal (0 Hz) while retaining Delta (0.5 - 4 Hz), Theta (4 – 8 Hz), Alpha (8 – 13 Hz), and Beta (13 – 32 Hz) bands. A fifth-order digital Butterworth filter was used for this process, chosen for its “maximally flat” response in the passband, which minimizes ripple effects despite a slower transition band (Shouran and Elgamli, 2020).
- **Normalization and segmentation**: All the signals were normalized to a zero mean and a standard deviation of one, reducing variability across databases and enhancing numerical stability. After this, signals were segmented based on sleep stages annotated by clinicians from hypnograms, resulting in four separate databases representing each sleep stage: N1, N2, N3, and REM.
- **Artifact removal**: Segments containing artifacts were removed based on annotations provided by experts for each session and their respective timestamps.

### 2.9. Signal features extraction

EEG signals were analyzed to extract seven features from time-domain and nonlinear features subsets, employing various signal-processing techniques and mathematical models (Bezbochina et al., 2023; Thakor and Tong, 2004; Alvarez et al., 2013; He et al., 2018; Álvarez et al., 2012).

For each signal segment annotated as a different sleep stage, the parameters were assessed as the mean of two-minute windows with a 50% overlap.

The final set of features was computed as the mean of the segments of each sleep stage (N1, N2, N3, and REM) for the whole overnight recording and through all EEG channels.

- **Time-domain features** provide insights into signal characteristics such as amplitude, variability, and distribution over time, helping to identify physiological dynamics and anomalies. Variance (indicating data distribution around the mean) and skewness measured signal dispersion and asymmetry, while kurtosis and maximum value represented peakedness and amplitude extremes (Thakor and Tong, 2004; Alvarez et al., 2013; Álvarez et al., 2012; He et al., 2018).
- **Non-linear parameters** analysis examines complex interactions within electrophysiological signals, highlighting their chaotic behavior and non-stationarity (Álvarez et al., 2012; Thakor and Tong, 2004). Sample Entropy quantifies complexity based on sequence similarity probability, where lower values indicate regularity and higher values reflect complexity. Lempel-Ziv complexity measures data intricacy through compression-based entropy, with higher values denoting increased complexity (Álvarez et al., 2012; Alvarez et al., 2013; Bezbochina et al., 2023).

This approach aimed to extract informative characteristics that describe signal dynamics and correlations (Bezbochina et al., 2023; Thakor and Tong, 2004). Appendix A provides detailed definitions and corresponding mathematical formulas for the time-domain and non-linear features extracted from EEG signals.

## 3. Statistical analysis

Descriptive statistics were obtained for various variables of relevance, defining the demographic and clinical characteristics of the patient group. Depending on the nature of the variable under examination, mean, median, standard deviation, or frequencies are estimated. The listwise deletion method was used to cope with missing data.

The associations between CSF biomarkers (GSA, AASA, CEL, CML, MDAL) and overall EEG features of both-sided central, frontal, and occipital signals and during N1, N2, N3, and REM sleep stages have been assessed using Spearman’s rank test. We also calculated adjusted correlations controlling for age, gender, and ApoE4 status. All tests were two-tailed, and p-values<0.05 were considered statistically significant. Initially, we investigated the interrelationships among CSF biomarkers (GSA, AASA, CEL, CML, MDAL). Correlation matrices were utilized for this analysis.

The study uncovered a substantial correlation among the targeted CSF biomarkers for prediction, indicating that predictions made independently would likely lack meaningful coherence. Hence, a combined approach to predict all CSF biomarkers is deemed appropriate. Given the constraints of the limited database, this insight is expected to significantly enhance the predictions’ accuracy.

Following that, we conducted EEG feature selection, retaining those features that showed a significant correlation (both unadjusted and adjusted) with at least one CSF biomarker. Descriptive statistics and Spearman’s test were carried out using Python (version 3.10.12).

## 4. Machine Learning methods for the prediction of CSF oxidative damage biomarkers

Subsequently, various methods were explored to predict CSF levels from the retained EEG features. These methods were performed using Python (version 3.10.12). The libraries employed were Scikit-Learn for the Random Forest implementation and Pyro (Bingham et al., 2018) for the Gaussian Process model.

### 4.1. Dimensionality reduction

To address the interdependencies among CSF biomarkers (Figure 3), we integrated them using Principal Component Analysis (PCA), a statistical technique that reduces the dimensionality of data while retaining as much variability as possible (Jolliffe and Cadima, 2016). For our analysis, we extracted one principal component (1PC), capturing 54.89% of the variance, with each CSF biomarker contributing with distinct weights.

Similarly, EEG data preprocessing involved handling missing values using Multiple Imputation by Chained Equations (MICE), which imputes values iteratively based on inter-variable relationships (Wulff and Jeppesen, 2017; Mera-Gaona et al., 2021). EEG features were also combined through PCA to reduce dimensionality and noise.

Using this biomarker and EEG components, models were trained for prediction, and predicted components were transformed back to their original scales via inverse matrix operations, yielding final values for each CSF biomarker.

### 4.2. Machine Learning approaches

To address this regression problem, we utilized two machine learning techniques: Random Forest (RF) and Gaussian Process (GP), each applied to the previously pre-processed database. The RF model, a supervised ensemble learning method, combines multiple decision trees trained independently to create a robust predictive framework (Sagi and Rokach, 2020). It was employed to predict principal component (1PC) derived from the CSF biomarker data, or each of the biomarkers separately (Bioms). This approach was implemented both with all EEG features extracted and transformed through Principal Component Analysis (PCA) and with the retained EEG without dimensionality reduction, enabling comparison of the model’s performance across different input representations.

In addition to its use for regression, the RF model was also applied to a binary classification task. By incorporating EEG data from healthy individuals into the dataset, we evaluated whether the same features used for biomarker prediction could reliably distinguish between AD patients and healthy controls. This classification analysis served as an additional validation step to confirm that the selected features reflect meaningful disease-related patterns rather than capturing spurious correlations.

On the other hand, the GP models a probability distribution over possible functions that fit a given set of data points, incorporating uncertainty into predictions through a covariance (kernel) function (Rasmussen, 2004). This method was also used to predict both 1PC and the biomarkers independently, enabling direct performance comparisons with RF and providing an additional layer of interpretability due to its ability to measure uncertainty.

To assess the contribution of individual EEG features to model performance, we computed feature importance scores using algorithm-specific approaches. In RF models, feature importance was estimated via the Mean Decrease in Impurity (MDI) method (Breiman, 2001). At each decision tree split, the reduction in impurity (measured as the decrease in residual variance) is attributed to the splitting feature. The final importance score of each feature is computed as the sum of these reductions across all trees, with higher scores indicating features more frequently used in informative splits (Breiman, 2001).

In GP models, we employed Automatic Relevance Determination (ARD) within the Radial Basis Function (RBF) kernel (Neal, 1996; Rasmussen and Williams, 2006). This method assigns a separate length-scale parameter to each input feature. Feature relevance was derived by taking the inverse of the learned length-scale, under the rationale that features with shorter length-scales have greater influence on model predictions(Neal, 1996; Rasmussen and Williams, 2006).

All importance values were normalized to allow for comparability across models.

## 5. Results

### 5.1. Characteristics of the cohorts

We initially recruited a consecutive series of 144 individuals with a recent diagnosis of mild to moderate AD. Out of these, 128 patients completed a full PSG (Gaeta et al., 2020). However, 70 cases were removed because their PSG recordings were no longer available, while several had missing data for some of the CSF biomarkers, resulting in a final sample of 42 patients. Table 1 lists the sample characteristics, which include socio-demographic information, comorbidities, cognitive assessments, CFS biomarker levels, APOE genotyping, self-reported drowsiness, and a summary of extracted polysomno-graphic parameters.

**Table 1:** Characteristics of patients with mild to moderate Alzheimer’s disease (AD). Key: AHI, apnea-hypopnea index; AI, arousal index; A*β*42, amyloid-beta protein; AASA, aminoadipic semialdehyde; CEL N*ε*-carboxyethyl-lysine; CML, N*ε*-Carboxymethyl-lysine; BMI, body mass index; ESS: Epworth sleepiness scale; IQR, interquartile range; p-tau, phospho-tau protein; SD, standard deviation of mean; t-tau, total-tau protein. GSA, glutamic semialdehyde; MDAL, N*ε*-malondialdehyde-lysine; MMSE, Mini-Mental State Examination; TIB, time in bed; t-tau, total-tau protein; TST, total sleep time; T90, cumulative sleep time percentage with oxyhemoglobin saturation (SpO2) < 90%.

| Data | number (%) or mean [SD] |
| --- | --- |
| **Sociodemographic data** |  |
| Women | 21 (50%) |
| Age, years | 74.71 [69.32; 80.11] |
| BMI, kg/m<sup>2</sup> | 27.96 [22.61; 33.30] |
| **Comorbidities** |  |
| Hypertension | 24 (57.14%) |
| Diabetes mellitus | 7 (16.67%) |
| Dyslipidemia | 21 (50%) |
| Heart diseases | 5 (11.90%) |
| Stroke | 2 (4.76%) |
| **Cognition** |  |
| MMSE | 23.31 [20.96; 25.66] |
| **Sleep-related symptoms** |  |
| ESS | 5.98 [2.27; 9.68] |
| Snoring | 1.36 [0.15; 2.56] |
| Not restorative sleep | 0.17 [-0.32; 0.66] |
| Witnessed apneas | 0.19 [-0.32; 0.70] |
| Insomnia | 0.33 [-0.24; 0.91] |
| **Polysomnographic parameters** |  |
| TIB, minutes | 412.98 [379.02; 446.94] |
| TST, minutes | 262.28 [170.2; 354.36] |
| Sleep efficiency, % | 63.67 [41.53; 85.8] |
| N1 sleep stage, % | 13.27 [5.40; 21.13] |
| N2 sleep stage, % | 25.96 [10.63; 41.29] |
| N3 sleep stage, % | 16.30 [4.32; 28.28] |
| REM sleep, % | 8.05 [1.05; 15.05] |
| Latency to N1, minutes | 36.95 [-12.45; 86.35] |
| Latency to REM sleep, minutes | 156.34 [58.04; 254.65] |
| AHI events/hour | 21.90 [5.30; 38.49] |
| AI events/hour | 38.91 [19.38; 58.44] |
| Average SpO<sub>2</sub>, % | 92.74 [90.39; 95.09] |
| T90% | 7.95 [-5.56; 21.46] |
| ODI events/hour | 28.89 [5.96; 51.83] |
| **AD core biomarkers** |  |
| Aβ42, pg/ml | 510.64 [376.79; 644.50] |
| t-Tau, pg/ml | 550.55 [281.51; 819.59] |
| p-Tau, pg/ml | 81.27 [48.57; 113.96] |
| **Oxidative damage biomarkers** |  |
| GSA, μmol/mol of lysine | 6556.15 [3884.08; 9228.23] |
| AASA, μmol/mol of lysine | 88.58 [50.61; 126.54] |
| CEL, μmol/mol of lysine | 116.07 [62.44; 169.71] |
| CML, μmol/mol of lysine | 1126.60 [813.04; 1440.17] |
| MDAL, μmol/mol of lysine | 210.68 [-21.64; 443.00] |

### 5.2. EEG features

In the Supplementary Material, Table I indicates the average values of the 114 EEG signal features for each EEG channel according to sleep phases. In addition, appendix B offers a list of the overall EEG features computed (Table B.3).

### 5.3. Correlations between EEG features and CSF biomarkers

We examined the relationships between qEEG features and CSF biomarkers. Figure 2 presents the statistically significant correlations, both before and after adjustment for potential confounding factors.

**Figure 2:**
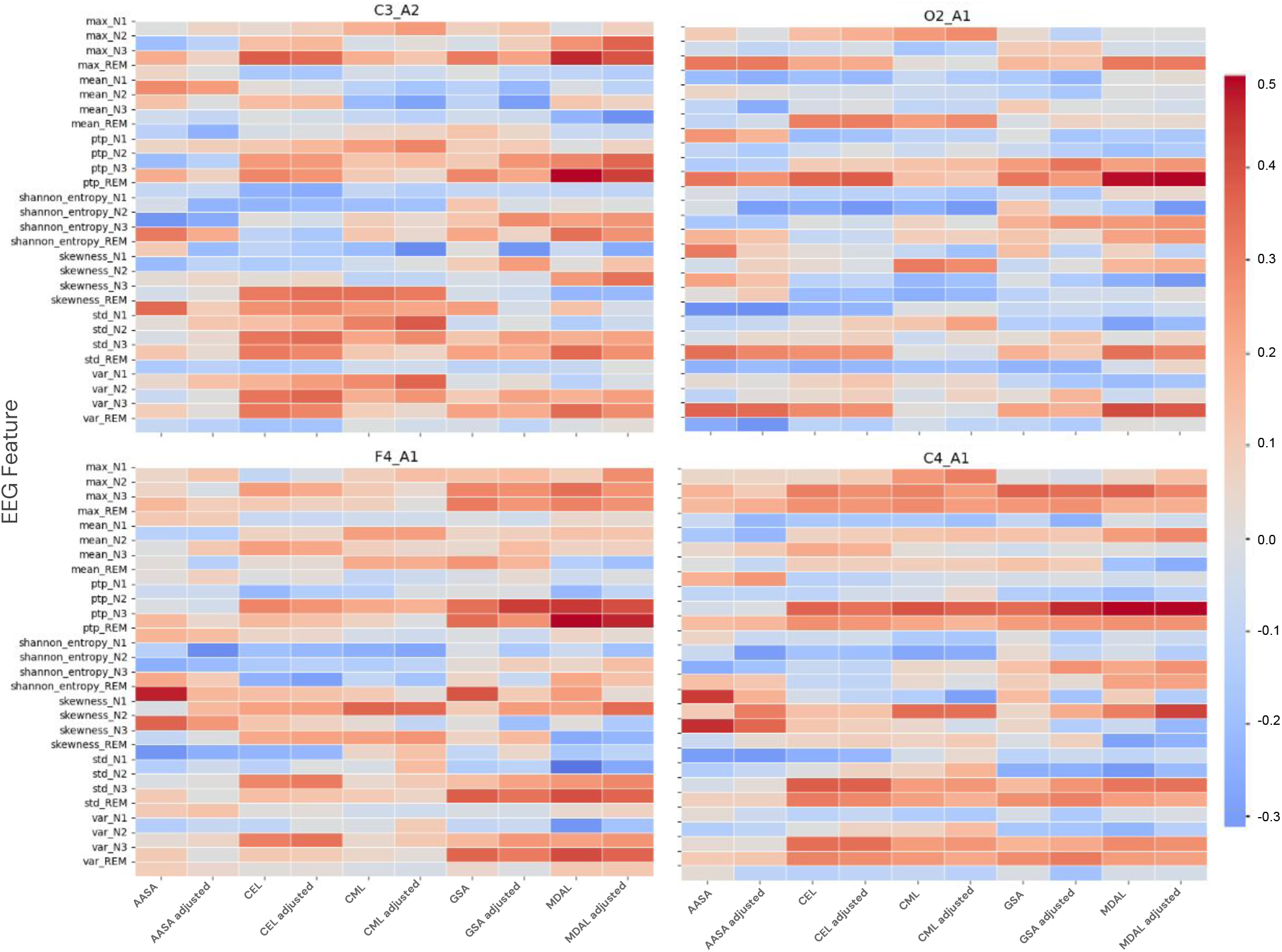
Significant correlations between sleep EEG features and cerebrospinal fluid biomarkers according to sleep stages for each one of the electrodes used. The represented values are unadjusted (Rho) or adjusted for age, sex, and ApoE4 status (Rho adjusted). Warmer colours indicate stronger positive correlations, while cooler colours represent weaker or negative correlations. Asterisks (*) highlight statistically significant correlations (p<0.05). Key: AASA, aminoadipic semialdehyde; CEL, N*ε*-carboxyethyl-lysine; CML, N*ε*-carboxymethyl-lysine; GSA, glutamic semialdehyde; MDAL, N*ε*-malondialdehyde-lysine; C3_A2, C3-A2 EEG channel; C4_A1, C4-A1 EEG channel; EEG, electroencephalogram; F3_A2, F3-A2 EEG channel; F4_A1, F4-A1 EEG channel; O1_A2, O1-A2 EEG channel; O2_A1, O2-A1 EEG channel; LempelZiv, Lempel-Ziv; max, maximum value; SampEnt, Sample Entropy; ShanEnt, Shannon Entropy.

Based on predefined selection criteria, the following EEG features were retained for analysis:

- Central region (C3, C4): Maximum amplitude and peak-to-peak amplitude during N2 and N3 sleep stages; variance, standard deviation, and skewness during N1 and N2; and Shannon entropy during REM sleep.
- Frontal region (F4): Peak-to-peak amplitude during N2 and N3; standard deviation during N1 and N3; variance during N3; and Shannon entropy during REM sleep.
- Occipital region (O2): Peak-to-peak amplitude, standard deviation, and variance during N3.

In addition, we examined the correlations among the CSF biomarkers themselves. Figure 3 illustrates these intercorrelations, highlighting strong associations across all biomarkers. This supports their integration into one principal component (1PC) to enhance predictive modeling accuracy.

**Figure 3:**
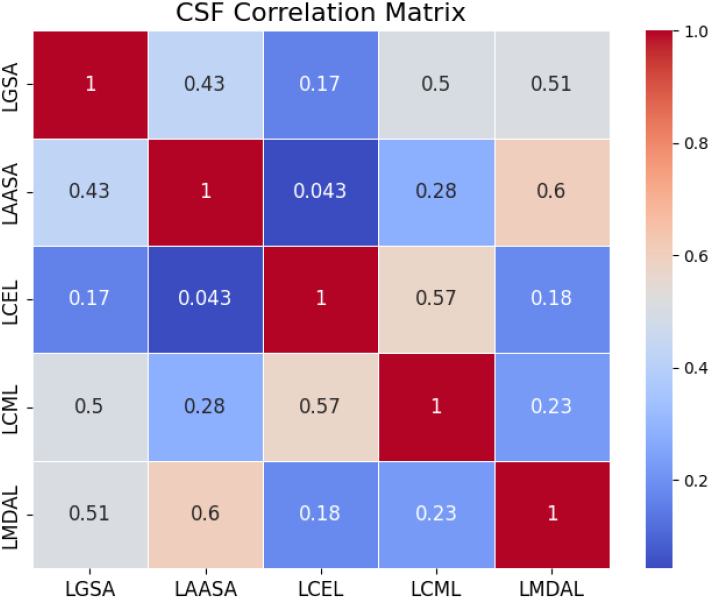
Heatmap and correlation matrix of cerebrospinal fluid (CSF) oxidative stress biomarkers. The matrix illustrates the pairwise Pearson correlation coefficients among glutamic semialdehyde (GSA), aminoadipic semi-aldehyde (AASA), N*ϵ*-carboxyethyl-lysine (CEL), N*ϵ*-carboxymethyl-lysine (CML), and N*ε*-malondialdehyde-lysine (MDAL). Warmer colours indicate stronger positive correlations, while cooler colours represent weaker or negative correlations. The observed intercorrelations suggest shared pathological mechanisms and justify the integration of these biomarkers into the principal component analysis (PCA) to reduce dimensionality and enhance predictive modeling accuracy in Alzheimer’s disease (AD) research.

### 5.4. CSF prediction

The performance metrics used to evaluate these models include R-squared (R^2^), Root Mean Squared Error (RMSE), Median Absolute Error (MeadAE), Mean Absolute Percentage Error (MAPE), Mean Absolute Error (MAE) as reported in Table 2. These metrics collectively provide a comprehensive view of model fit and prediction accuracy.

**Table 2:**
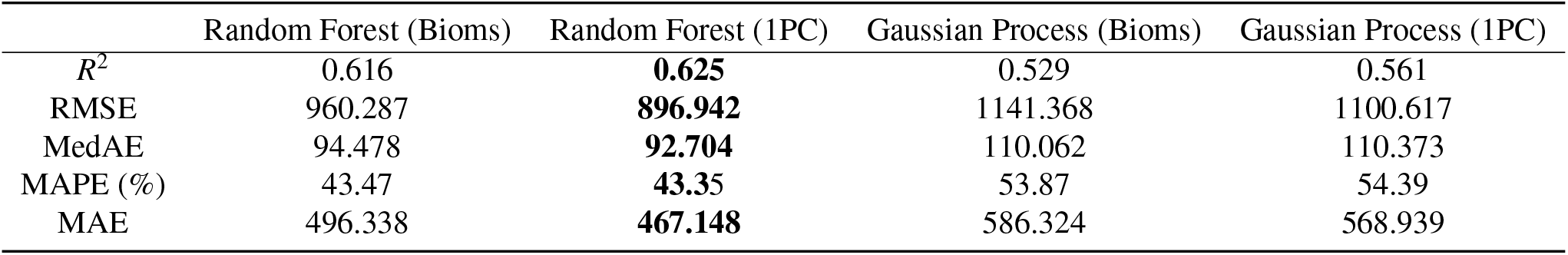
Comparison of performance metrics for CSF biomarker prediction across different models. The table reports the mean test set values for five metrics: *R*^2^ score, RMSE, MedAE, MAPE, and MAE. Results are shown for four models: RF with separate predictions for each biomarker, RF with 1PC, GP with a separate prediction, and GP with 1PC. Higher *R*^2^ values indicate better model fit, while lower RMSE, MedAE, MAPE, and MAE values indicate greater accuracy. In bold, the best-performing results are highlighted. RMSE, Root Mean Squared Error; *R*^2^, R-squared; MeadAE, Median Absolute Error; MAPE, Mean Absolute Percentage Error; MAE, Mean Absolute Error.

|  | Random Forest (Bioms) | Random Forest (1PC) | Gaussian Process (Bioms) | Gaussian Process (1PC) |
| --- | --- | --- | --- | --- |
| $R^2$ | 0.616 | **0.625** | 0.529 | 0.561 |
| RMSE | 960.287 | **896.942** | 1141.368 | 1100.617 |
| MedAE | 94.478 | **92.704** | 110.062 | 110.373 |
| MAPE (%) | 43.47 | **43.35** | 53.87 | 54.39 |
| MAE | 496.338 | **467.148** | 586.324 | 568.939 |

R^2^ quantifies how well the regression model explains the variance in the dependent variable, indicating the proportion of variability accounted for by the predictor variables (Chicco et al., 2021). Its values range from 0 to 1, with higher values indicating better model performance. RMSE measures the square root of the average squared difference between predicted and actual values, providing a sensitive indicator of large errors. MedAE reflects the median of absolute errors, offering robustness to outliers. MAPE expresses prediction error as a percentage, making it interpretable across scales, while MAE represents the mean of the absolute errors, giving a linear view of prediction deviations. Lower values of RMSE, MedAE, MAPE, and MAE indicate more accurate predictions (Botchkarev, 2018).

Among the models tested, the RF trained to predict a single principal component (1PC) of CSF biomarkers yielded the most favorable overall results. It achieved the highest mean test set *R*^2^ score (0.625), indicating the best model fit. It also outperformed all other models across the remaining four metrics. The RF model with separate predictions for each biomarker followed closely in performance, particularly in terms of RMSE (960.29) and MAE (496.34).

It is important to highlight that the reported performance metrics reflect average values across test set patients. As illustrated in the individual prediction visualizations (Figure 5), there is an inter-patient variability. However, the goal of the model was not to precisely predict the exact biomarker values, but rather to generate reliable approximations. This objective is met, as seen in the per-patient, per-biomarker predictions, which demonstrate consistently stable trends and error distributions.

A second model, trained by merging twelve principal components for merging all EEG features (capturing 87.26% of the database variance) and 1PC for biomarkers, achieved similar overall metrics and errors (data not shown). However, its consistency across biomarkers was slightly reduced, suggesting that direct training with retained EEG variables may offer more robust predictions.

Additionally, several EEG features consistently emerged as top predictors across both RF and GP models, highlighting their relevance in capturing neurophysiological patterns linked to CSF biomarker levels. As shown in Figure 4, we focus on the top 8 out of the 15 most informative features. This threshold was set pragmatically based on the observed drop in importance values through all models, allowing us to concentrate the analysis on the most relevant contributors. These key features include:

**Figure 4:**
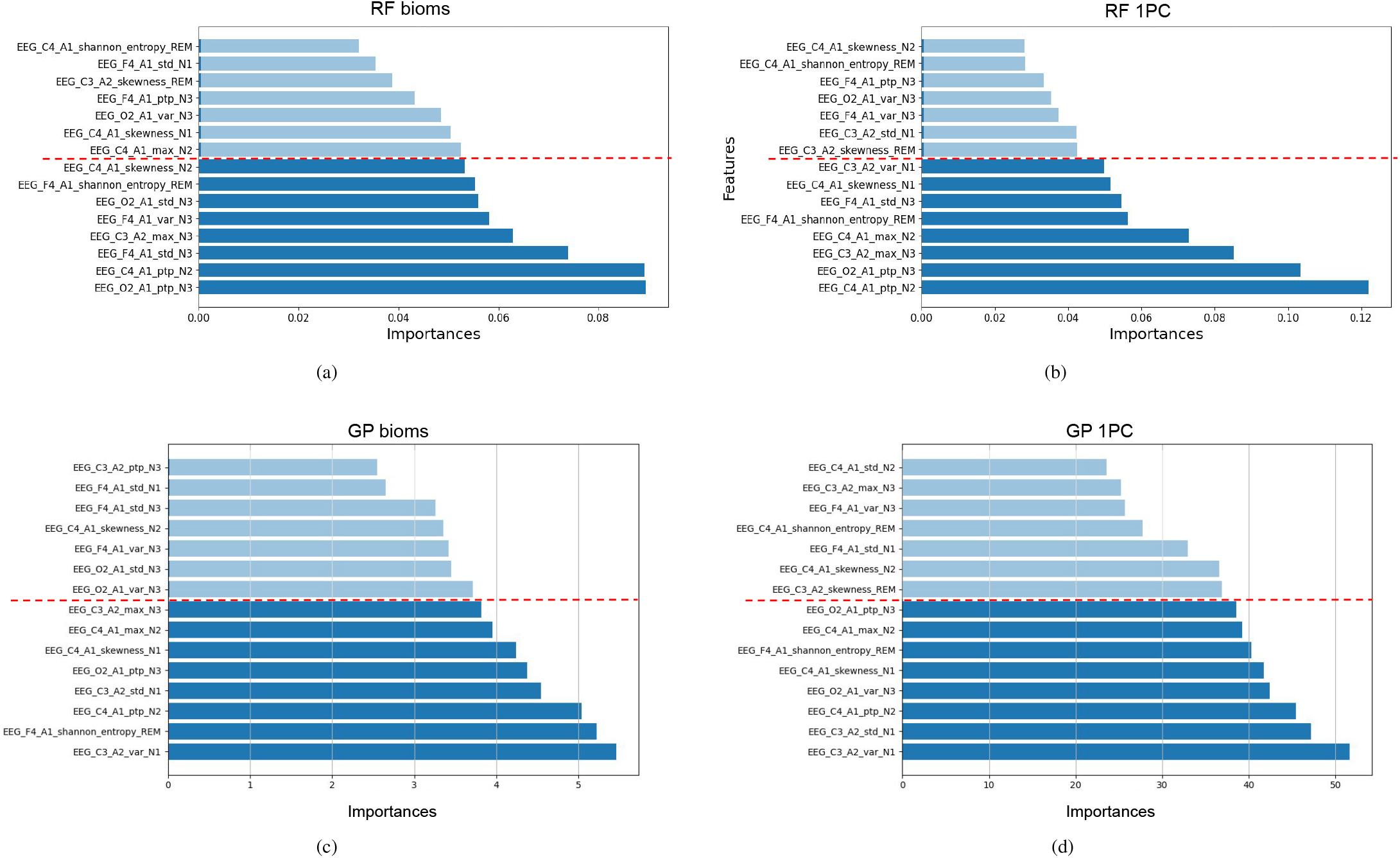
Feature importance scores for the prediction of cerebrospinal fluid oxidative stress biomarkers using sleep qEEG-derived features. Panels (a) and (b) show results from Random Forest models (RF) trained to predict individual biomarkers (a) or their first Principal Component (1PC) (b). Panels (c) and (d) show corresponding results for Gaussian Process (GP) models. The eight most relevant features for each model are highlighted. The most predictive features were predominantly derived from non-REM sleep stages (N1-N3), particularly from central and occipital EEG channels. Key features included peak-to-peak amplitude, variance, skewness, and Shannon entropy, underscoring the contribution of time-domain and nonlinear EEG characteristics to the modeling of oxidative stress-related alterations in early Alzheimer’s disease.

- *EEG***_***O*2**_***A*1**_***ptp***_***N*3 (peak-to-peak amplitude, occipital region, N3 sleep) – important in all models,
- *EEG***_***C*4**_***A*1**_***ptp***_***N*2 (peak-to-peak amplitude, central region, N2 sleep) – top-ranked across all models,
- *EEG***_***F*4**_***A*1**_***shannon***_***entropy***_***REM* (Shannon entropy, frontal region, REM) – consistently informative all models,
- *EEG***_***C*4**_***A*1**_***skewness***_***N*2*/N*1 (skewness, central region, N1–N2) – significant in both RF and GP 1PC settings,
- *EEG***_***C*3**_***A*2**_***var***_***N*1 and *EEG***_***C*3**_***A*2**_***std***_***N*1 (variance and standard deviation, central region, N1 sleep) – dominant in GP models.

Moreover, within each individual model, the majority of top-ranked predictors originate from SWS, underscoring this stage as the most informative and consistently contributing the greatest number of key features.

A key advantage of GP modeling is its ability to estimate uncertainty. However, the associated error margins are relatively large, especially at higher biomarker levels, indicating lower confidence in predictions. This may stem from the limited dataset size, particularly the scarcity of samples from patients with advanced AD.

### 5.5. Validation with healthy controls

To assess the robustness and disease-specificity of our regression models, we performed additional validation analyses using a control cohort of cognitively healthy individuals. Importantly, this group lacked measured CSF biomarker values and, thus, was excluded from model training. Nonetheless, the same EEG features extracted from their sleep recordings were used as input to the previously trained models to evaluate generalization and specificity.

Predicted biomarker values for healthy controls were consistently distinct from both the true and predicted values of AD patients, typically falling outside the range of AD distribution. While the model does not fully capture the extremes of the AD biomarker spectrum—likely due to limited training data—it still produces meaningful and biologically relevant predictions that differentiate between groups Figure 5.

**Figure 5:**
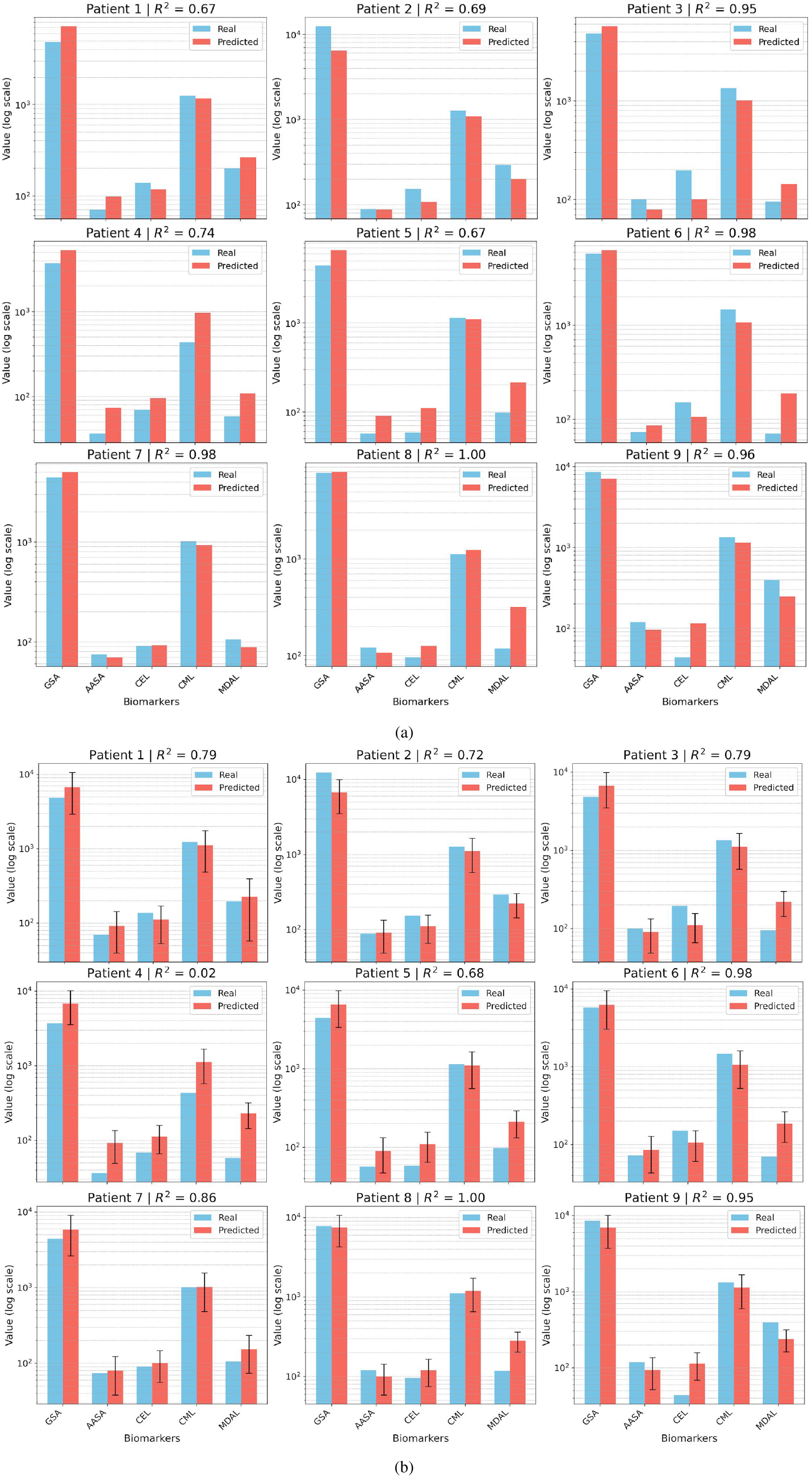
R^2^ scores for test predictions across all biomarkers, shown per patient on a logarithmic scale to enhance visualization. Results are displayed for the two models trained to predict a single principal component (1PC): (a) Random Forest and (b) Gaussian Process, incorporating uncertainty values associated with each prediction. Each plot compares predicted vs. true values for cerebrospinal fluid (CSF) oxidative stress biomarkers. Key: AASA, aminoadipic semialdehyde; CEL, Nε-carboxyethyl-lysine; CML, Nε-carboxymethyl-lysine; GSA, glutamic semialdehyde; MDAL, Nε-malondialdehyde-lysine; N3, N3 sleep stage; max, maximum value; R^2^, R-squared; ShanEnt, Shannon Entropy.

To further validate that EEG features reflect disease-related changes, we trained a Random Forest classifier to distinguish AD patients from healthy controls using the same EEG inputs (data not shown). The model achieved strong performance (*AUC* = 0.99, *Accuracy* = 0.91), reinforcing the idea that the selected features encode pathophysiologically meaningful signals rather than random variance.

## 6. Discussion

In this study, we developed advanced ML models to evaluate the predictive capacity of nonlinear and time-domain features derived from sleep EEG for estimating CSF protein oxidation markers in early-stage AD. Our results show that peak-to-peak amplitude, variance, skewness, and standard deviation during NREM sleep, as well as Shannon entropy during REM sleep, particularly in central and frontal brain regions, are strong predictors of non-enzymatic oxidative stress biomarkers. These biomarkers, including protein oxidation, AGEs, and ALEs, are well-established indicators of irreversible oxidative damage in the AD brain.

### 6.1. Integration of Functional and Molecular Markers

Our results introduce a novel, clinically relevant framework that integrates functional neurophysiological data (sleep EEG) with molecular biomarkers, bridging two fundamental aspects of AD pathology: network dysfunction and oxidative molecular damage (Tönnies and Trushina, 2017; Czekus et al., 2022). By leveraging non-invasive sleep EEG alongside CSF biomarker analysis, we enhance the biological specificity of qEEG as a tool for early detection.

Recent studies indicate that nonlinear EEG metrics detect AD-related brain alterations that traditional frequency-based analyses often miss (Jeong, 2004; Chetty et al., 2024). Features such as Shannon Entropy capture declining brain signal complexity, while variance and skewness reflect regional connectivity disruptions, characteristics of AD (Karaca and Moonis, 2022; Aoki et al., 2023; Koenig et al., 2005). However, limited research has explored these markers specifically in sleep EEG in AD; a gap this study addresses.

### 6.2. Bidirectional Relationship Between Sleep and Oxidative Stress

Our findings, indicating that SWS and REM sleep yield the most informative qEEG signals, reinforce prior evidence of a bidirectional interplay between sleep disturbances and oxidative stress. (Czekus et al., 2022). Disruptions in SWS and REM architecture not only reflect the impact of oxidative damage on thalamocortical and corticocortical circuits involved in sleep regulation but also drive further reactive oxygen species production (Mander et al., 2016; Lucey et al., 2019; Mathangi et al., 2012).

SWS plays a key role in cellular repair, mitochondrial function, and antioxidant defence (Mander et al., 2016). Reduced SWS duration or fragmentation is associated with elevated CSF oxidative stress markers and accelerated AD progression (Lucey et al., 2019). Similarly, the glymphatic system, which clears neurotoxic waste such as A*β*, operates most effectively during NREM sleep, particularly SWS (Hablitz et al., 2019; Xie et al., 2013). Disruptions to A*β* clearance from SWS impairment or hypoxia (e.g., sleep apnea) may promote A*β* accumulation and neuronal vulnerability (Kyrtsos and Baras, 2015).

REM sleep also plays a critical role in neurotoxin clearance and antioxidant regulation. Its deprivation increases oxidative stress, impairs cognitive function, and depletes antioxidants, linking reduced REM sleep to lower CSF A*β* levels and cholinergic neurons degeneration in AD (Mathangi et al., 2012; Mallick and Gulyani, 1996; Ramanathan et al., 2002; Mander et al., 2016).

Our findings align with region-specific oxidative damage in AD, showing that qEEG features predictive of oxidative stress markers are most prominent in the frontal brain during SWS and REM sleep (Romanella et al., 2021; Chylinski et al., 2022; Takeda et al., 2001).

### 6.3. Methodological Considerations in the Use of Supervised Learning Models

The application of supervised learning models such as RF and GP offers complementary advantages and methodological challenges in the context of biomarker prediction from EEG data. One key strength of GP modeling is its ability to estimate predictive uncertainty, which allows the model to express confidence levels in its output. However, the observed increase in error margins, particularly at higher biomarker levels, raises concerns regarding the model’s reliability in extreme cases. This limitation is likely influenced by the relatively small sample size and the underrepresentation of patients with advanced AD in the training data.

Although RF models do not natively provide uncertainty estimates, the parallel observation of increased prediction error at higher biomarker values suggests that both approaches are affected by a common challenge: accurately modeling the distributional extremes of CSF biomarkers. This pattern has important implications for clinical translation, where extreme values may be the most diagnostically informative yet also the most difficult to predict.

To further explore model generalization and disease specificity, we assessed the performance of both regression and classification models on a control cohort of cognitively healthy individuals. These analyses, while not contributing to training, demonstrated the capacity of the EEG-based models to differentiate between pathological and non-pathological signal patterns, even in the absence of true biomarker values. This suggests that the EEG features extracted retain disease-related information and that the supervised models are able to exploit these signals meaningfully.

### 6.4. Strengths and Limitations

Our study has several key strengths. First, we analyzed a rigorously defined cohort of mild to moderate AD patients, ensuring a well-characterized population. Second, we utilized a comprehensive assessment battery, including overnight PSG and CSF biomarker analysis, a combination rarely used in AD research. Third, we included only drug-free participants to eliminate potential medication-induced distortions in REM sleep EEG features (Azami et al., 2023).

A major strength of our approach is the integration of ML to uncover complex, nonlinear relationships between qEEG features and CSF oxidative stress biomarkers, relationships that traditional linear methods often overlook (Chang et al., 2021; Lovejoy et al., 2021). Unlike prior AD studies that primarily relied on linear EEG metrics (D’Atri et al., 2021; Mander et al., 2016), our work incorporates nonlinear and time-domain EEG features, offering a deeper understanding of oxidative stress-related neurophysiological alterations.

Despite these strengths, some limitations must be acknowledged. The primary constraint is the small dataset from a single center, which may limit generalizability. However, our use of advanced ML models helped mitigate this issue by enhancing predictive performance. While a low number of patients with high CSF biomarker levels contributed to increased prediction error, especially at extreme biomarker values, both RF and GP models reliably captured trends in the data, consistently identifying key EEG features (see Figure 4). Furthermore, GP modeling provided uncertainty estimates that confirm stronger confidence in lower-to-mid range predictions, which could improve with larger datasets.

Other limitations include the use of only four EEG channels, which restricts detailed topographic analysis. The lack of CSF data in the healthy cohort, due to ethical concerns surrounding CSF collection, precluded direct evaluation of prediction error for controls. Nonetheless, the divergence in predicted biomarker distributions and the results of the classification task provide indirect but compelling support for the model’s validity. Additionally, while nonenzymatic protein biomarkers show promise for early AD risk assessment and disease progression, their clinical validation remains ongoing (Dakterzada et al., 2023).

### 6.5. Clinical and Research Implications

This integrative qEEG-CSF biomarker strategy could pioneer a paradigm shift in AD risk stratification and early diagnosis. By demonstrating that non-invasive sleep EEG features could estimate key oxidative stress biomarkers, we propose a cost-effective alternative to invasive CSF sampling (Monllor et al., 2021).

Moving away from invasive molecular markers toward a non-invasive, sleep-based monitoring system could enable clinicians to identify individuals at risk for AD—where early intervention is critical (Tan et al., 2014; Jack Jr et al., 2024)—as well as track disease progression in AD patients with heightened oxidative stress-related network damage.

This approach could be particularly valuable for large-scale screening of asymptomatic individuals, allowing for early intervention before clinical symptoms emerge. It can be seamlessly integrated into longitudinal monitoring protocols, complementing traditional biomarkers and neuroimaging methods (Jack Jr et al., 2024).

Additionally, sleep is a modifiable risk factor for oxidative stress, making this strategy highly relevant for personalized prevention. Targeted lifestyle and sleep interventions - including cognitive behavioral therapy for insomnia (CBT-I), transcranial stimulation, or melatonin therapy - could improve SWS and REM sleep, reduce oxidative damage, and potentially delay or prevent symptom onset (Romanella et al., 2021; Lucey, 2020; Dhapola et al., 2024).

Furthermore, EEG-derived features could serve as biomarkers to assess treatment efficacy, offering real-time insights into the effects of oxidative stress-targeting therapies on neural activity (Dhapola et al., 2024). Lastly, our methodology may propel research efforts toward the validation of CSF oxidatively modified protein biomarkers for clinical use without requiring invasive sampling procedures.

### 6.6. Future Directions

Future research should validate these findings in larger, diverse cohorts, including preclinical AD cases and healthy older adults, to distinguish AD pathology from normal aging (Jack Jr et al., 2024). Longitudinal studies are crucial for evaluating the stability and predictive value of qEEG biomarkers over time, as well as their role in disease progression and treatment response. Factors such as obstructive sleep apnea (OSA) and the ApoE genotype, which can exacerbate oxidative damage and potentially affect the accuracy of qEEG-based biomarkers, should be considered (Minakawa et al., 2019; Di Domenico et al., 2016; Xiong et al., 2021).

Future studies should explore how qEEG-based oxidative stress markers correlate with established AD biomarkers (*Aβ*, p-tau, t-tau) to refine risk models. High-density EEG should also be incorporated to improve spatial resolution, enabling a more detailed topographic analysis of AD-related network disruptions (Gordon and Rzempoluck, 2004; Stoyell et al., 2021). Lastly, intervention-based studies should evaluate whether sleep-targeted therapies (Dhapola et al., 2024) can modulate qEEG features, reduce oxidative stress, and delay cognitive decline.

On the other hand, from a methodological perspective, adequate representation of participants with extreme biomarker values—who remain underrepresented but are critical for improving model robustness and generalization—should be ensured. Furthermore, approaches that integrate uncertainty estimation (either inherently, as in GP models, or through post hoc calibration in tree-based methods) could enhance the interpretability and clinical trustworthiness of EEG-based predictive models. Lastly, AI-driven approaches could automate qEEG feature selection and classification, improving diagnostic accuracy while deep learning techniques enhance model generalization (Kabir et al., 2023; Gerla et al., 2019).

## 7. Conclusions

Our study demonstrates that advanced nonlinear and time-domain sleep EEG features, combined with an ML, reliably estimate CSF protein oxidation-derived markers in early AD.

This approach outperforms traditional methods and offers a non-invasive, cost-effective alternative to lumbar puncture sampling. Future research will focus on developing customized algorithms for different patient populations, refining predictive models, and integrating qEEG biomarkers into early AD detection frameworks. If validated, this strategy could facilitate personalized risk assessment and targeted interventions aimed at reducing oxidative stress and slowing neurodegeneration in aging populations.

## Data availability

The data supporting this study are available upon reasonable request to the corresponding author.

## CRediT authorship contribution statement

Anna Michela Gaeta: Conceptualization, Writing – original draft, Investigation, Methodology, Visualization. Lorena Gallego Viñarás: Formal analysis, Writing – original draft, Methodology, Investigation, Data curation, Visualization. Ferran Barbé: Project administration, Writing – review & editing, Investigation, Methodology. Pablo M. Olmos: Writing – review & editing, Formal analysis, Methodology, Investigation, Visualization. Reinald Pamplona: Methodology, Data curation, Writing – review & editing. Farida Dakterzada: Methodology, Data curation, Writing – review & editing. Arrate Muñoz-Barrutia: Project administration, Investigation, Formal analysis, Methodology, Writing – review & editing. Gerard Piñol-Ripoll: Project administration, Conceptualization, Investigation, Methodology, Writing – review & editing.

## Acknowledgements

We want to offer our heartfelt appreciation to all the patients and staff at the Hospital Universitari Santa Maria’s Sleep and Dementia Unit. The IRBLleida Biobank (B.0000682) and plataforma biobancos PT17/0015/0027/ also assisted us. All authors have revised and approved the submission of the manuscript. The authors utilized the language model ChatGPT, developed by OpenAI, to assist in drafting this paper.

## Appendix A. Definitions and mathematical formulas for quantitative EEG features

This supplementary material provides detailed definitions and corresponding mathematical formulas for the time-domain and non-linear features extracted from EEG signals.

### Appendix A.1. Time Domain Statistics

- *Variance* (*Var*) Variance measures the dispersion of the EEG signal values around the mean.

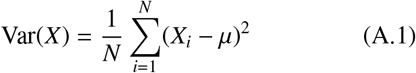

where µ is the mean of the signal *X*, and *N* is the number of samples (Alvarez et al., 2013).
- *Skewness (Skew)*
Skewness measures the asymmetry of the probability distribution of the EEG signal.

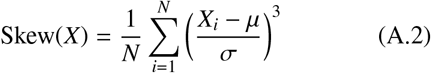

where σ is the standard deviation of the signal *X* (Alvarez et al., 2013).
- *Kurtosis (Kurt)* Kurtosis measures the tailedness of the probability distribution of the EEG signal (Alvarez et al., 2013).

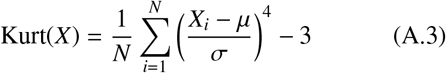
- *Mean* Mean amplitude of the EEG signal (Sulaiman et al., 2022).

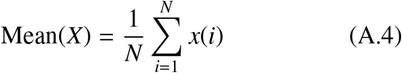
- *Peak-to-peak amplitude (PPAF)* PPAF quantifies fluctuations in EEG by measuring successive peak-to-peak amplitude differences, capturing regions of heightened electrical activity often associated with seizures (Bahhah and Attar, 2024).

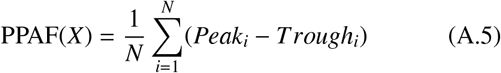

where *Peak*_*i*_ is the amplitude of the *i*-th peak in the EEG signal, *Trough*_*i*_ represents the amplitude of the *i*-th trough following the *i*-th peak and *N* the total number of peak-to-trough pairs.
- *Standard Deviation (SD)* Quantifies the variability or dispersion of the EEG signals. (Sulaiman et al., 2022).

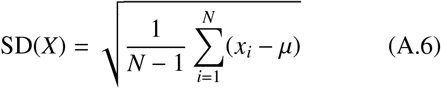

where µ is the mean.
- *Maximum Value (max)* The maximum value is the highest amplitude observed in the EEG signal (He et al., 2018).

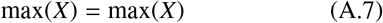

### Appendix A.2. Non-Linear Features

- *Shannon Entropy (ShanEnt)* Shannon Entropy quantifies the uncertainty or randomness in the EEG signal.

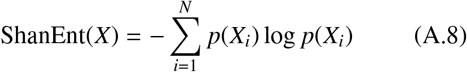

where *p*(*X*_*i*_) is the probability of the *i*-th signal value (Bezbochina et al., 2023).
- *Sample Entropy (SampEnt)* Sample Entropy measures the complexity and irregularity of the EEG signal.

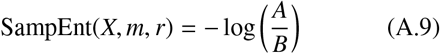

where *A* is the number of matching templates of length *m* + 1 and *B* is the number of matching templates of length *m* within a tolerance *r* (Alvarez et al., 2013).
- *Lempel-Ziv (LempZiv)* Lempel-Ziv Complexity measures the complexity of the EEG signal based on the number of distinct substrings and their recurrence rates. The formula is given by:

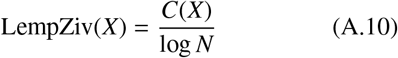

where *C*(*X*) is the number of distinct substrings in the signal *X* and *N* is the length of the signal (Alvarez et al., 2013).

## Appendix B. Quantitative EEG signals features

**Table B.3:**
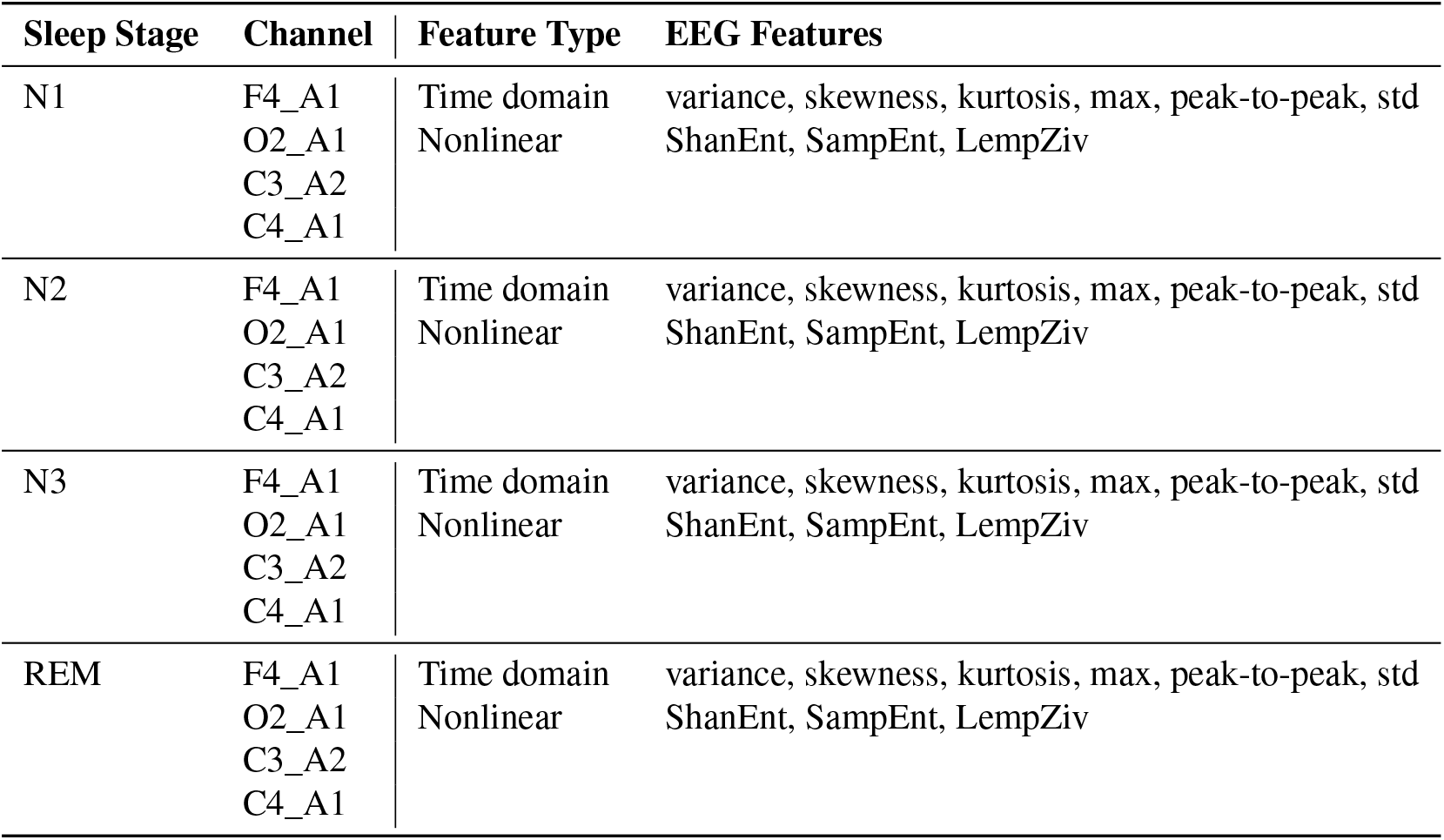
EEG features computed for the EEG F3-A2, F4_A1, C3_A2, C4_A1, O1_A2, O2_A1 channels, segmented by sleep stages. Key: EEG, Electroencephalogram; LempelZiv, Lempel Ziv complexity; max, maximum value; std; standard deviation, SampEnt, Sample Entropy; ShanEnt, Shannon Entropy. The notation used throughout the manuscript when referring to specific features follows the format: EEG_*’Channel’*_*’EEG Feature’*_*’Sleep Stage’*.

