## Supplemental Table 1 for "Non-linear and time-domain sleep qEEG features predict CSF protein damage markers in early Alzheimer’s disease"

| Channel | Sleep Stage | Variance | Skewness | Max | Shannon Entropy | Peak-to-Peak |
|---|---|---|---|---|---|---|
| F4_A1 | N1 | 0.736 ± 0.188 | -0.272 ± 0.483 | 2.767 ± 0.450 | 2.886 ± 0.058 | 6.007 ± 0.943 |
| | N2 | 0.881 ± 0.198 | -0.294 ± 0.417 | 2.845 ± 0.413 | 2.882 ± 0.056 | 6.289 ± 0.889 |
| | N3 | 1.477 ± 0.423 | -0.294 ± 0.375 | 3.335 ± 0.532 | 2.857 ± 0.060 | 7.402 ± 1.110 |
| C3_A2 | REM | 0.588 ± 0.215 | -0.254 ± 0.520 | 2.369 ± 0.424 | 2.890 ± 0.060 | 5.174 ± 1.019 |
| | N1 | 0.786 ± 0.195 | 0.066 ± 0.295 | 3.134 ± 0.607 | 2.865 ± 0.027 | 6.158 ± 0.942 |
| | N2 | 0.936 ± 0.179 | 0.014 ± 0.228 | 3.218 ± 0.556 | 2.867 ± 0.022 | 6.427 ± 0.804 |
| C4_A1 | N3 | 1.531 ± 0.382 | -0.033 ± 0.125 | 3.757 ± 0.601 | 2.843 ± 0.021 | 7.539 ± 1.011 |
| | REM | 0.547 ± 0.167 | 0.132 ± 0.327 | 2.605 ± 0.646 | 2.859 ± 0.026 | 5.006 ± 0.934 |
| | N1 | 0.793 ± 0.223 | -0.173 ± 0.362 | 2.940 ± 0.414 | 2.869 ± 0.031 | 6.228 ± 0.929 |
| | N2 | 0.950 ± 0.228 | -0.155 ± 0.270 | 3.075 ± 0.389 | 2.868 ± 0.023 | 6.526 ± 0.850 |
| | N3 | 1.531 ± 0.434 | -0.138 ± 0.174 | 3.566 ± 0.549 | 2.844 ± 0.021 | 7.554 ± 1.164 |
| | REM | 0.582 ± 0.207 | -0.211 ± 0.393 | 2.382 ± 0.440 | 2.870 ± 0.027 | 5.209 ± 1.050 |
| O2_A1 | N1 | 0.867 ± 0.227 | -0.106 ± 0.461 | 3.007 ± 0.439 | 2.864 ± 0.032 | 6.255 ± 1.021 |
| | N2 | 0.884 ± 0.193 | -0.062 ± 0.377 | 3.002 ± 0.421 | 2.855 ± 0.024 | 6.192 ± 0.920 |
| | N3 | 1.241 ± 0.324 | -0.043 ± 0.279 | 3.339 ± 0.508 | 2.832 ± 0.020 | 6.827 ± 1.005 |
| | REM | 0.571 ± 0.205 | -0.177 ± 0.500 | 2.353 ± 0.428 | 2.866 ± 0.027 | 5.039 ± 1.017 |

Table 1: Mean EEG signal feature values grouped by EEG channel and sleep stages. Key: C3_A2, C3-A2 EEG channel; C4_A1, C4-A1 EEG channel; EEG, Electroencephalogram; F3_A2, F3-A2 EEG channel; F4_A1, F4-A1 EEG channel; LempelZiv, Lempel Ziv complexity; max, maximum value; O1_A2, O1-A2 EEG channel; O2_A1, O2-A1 EEG channel; p-tau, phospho-tau protein; SampEnt, Sample Entropy; ShanEnt, Shannon Entropy.

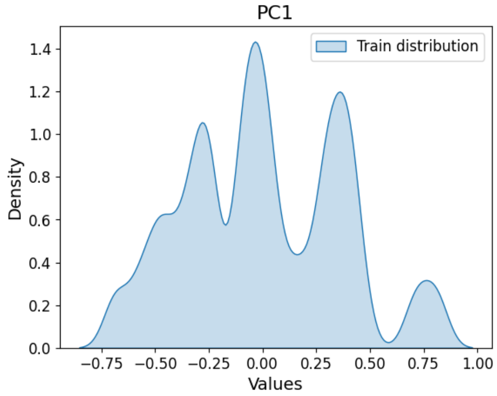


Figure 1: Distribution of Principal Component 1 (PC1) derived from CSF biomarker data. The distribution aligns with four distinct Gaussian curves, suggesting underlying subgroups within the dataset. This informed the Gaussian Process model, enabling more precise biomarker predictions by training separate models for each distribution segment.
